# Cluster-permutation statistical analysis for high-dimensional brain-wide functional connectivity mapping

**DOI:** 10.1101/849554

**Authors:** Jose M. Sanchez-Bornot, Maria E. Lopez, Ricardo Bruña, Fernando Maestu, Vahab Youssofzadeh, Su Yang, Paula L. McLean, Girijesh Prasad, KongFatt Wong-Lin

## Abstract

Brain functional connectivity (FC) analyses based on magnetoencephalographic (MEG) signals have yet to exploit the intrinsic high-dimensional information. Typically, these analyses are constrained to regions of interest to avoid the curse of dimensionality, which leads to conservative hypothesis testing. We removed such constraint by extending cluster-permutation statistics for high-dimensional MEG-FC analyses. We demonstrated the feasibility of this approach by identifying MEG-FC resting-state changes in mild cognitive impairment (MCI), a prodromal stage of Alzheimer’s disease. We found dense clusters of increased connectivity strength in MCI compared to healthy controls (hypersynchronization), in delta (1-4 Hz) and higher-theta (6-8 Hz) bands oscillations. These clusters mainly consisted of interactions between occipitofrontal and occipitotemporal regions in the left hemisphere and could potentially be used as neuromarkers of early progression in Alzheimer’s disease. Our novel approach can be used to generate high-resolution statistical FC maps for neuroimaging studies in general.

## Introduction

Functional connectivity (FC) analyses are continuously evolving, helping us to shape our understanding of network organization in healthy and unhealthy brains^1–4^. Typically, FC studies are conducted in resting-state since the associated spontaneous brain activity recruits multiple brain regions and networks, which can also be observed during active cognitive states^5–9^. Due to the consistency of resting-state FC results across multiple datasets, it also makes the study of brain disorders possible^2–5,10^. Furthermore, the use of resting-state functional magnetic resonance imaging (rs-fMRI) has attracted most attention given an excellent spatial resolution of fMRI to map brain function differences between conditions^6,7^. However, rs-fMRI analyses provide only an ultra-low frequency filtered and indirect representation of the underlying neural dynamics, as fMRI is based on the slow blood-oxygen-level dependent (BOLD) signal^11^. In contrast, electro/magnetoencephalography (EEG/MEG) imaging resolves such limitations by directly reflecting transient neural dynamics and allowing to infer communication among brain regions^12^.

In any case, either using fMRI^7–9^ or EEG/MEG^13–16^ data, analyses are heavily reliant on the use of regions of interest (ROIs) for reducing dimensionality, with a trade-off between the advantages of faster computations, simpler and less-conservative statistical tests, versus the possible loss of information and biased results^16,17^. Conversely, FC studies in the last decade have shown the feasibility of high-dimensional approaches to study network dynamics in greater detail^12,17–23^; e.g. using cluster-permutation statistics, with the critical advantage that significant network clusters ensure strong evidence of inter-regional connectivity^17–19,22^. Therefore, high-dimensional FC analysis could enhance the evaluation of FC differences between healthy and unhealthy brain conditions^19,22^. However, the state-of-the-art applications of cluster-permutation statistics are mostly limited to non-EEG/MEG data^17–19,21,22^ and low-dimensional analysis^23^, and hence not fully exploiting the advantages of the latter approach.

In this work, we extended the application of cluster-permutation statistics to truly high-dimensional scenarios, e.g. using a source-based FC analysis after solving the EEG/MEG inverse problem. Specifically, we estimated source pairwise FC using our recently proposed connectivity measure to control for volume conduction effects^24^. By concurrently dealing with FC analysis in the intrinsic high-dimensional space while controlling for volume conduction, we demonstrated increased sensitivity of post-hoc statistical analyses. This approach was applied to a dataset of 30 healthy control (HC) and 30 mild cognitive impairment (MCI) participants, where MCI was diagnosed according to standard criteria^25^. Statistical tests for the estimated MEG-FC networks differences between the HC and MCI groups, and for the covariation of these networks with respect to measured cognitive tests, were evaluated. We found significantly increased activation of occipitotemporal and occipitofrontal networks in MCI with respect to HC participants (hypersynchronization) in the left hemisphere, possibly associated with cognitive decline, and showed that significant source MEG-FC clusters can potentially be used for biomarker research in Alzheimer’s disease (AD).

## Results

We will focus on source MEG-FC analysis to demonstrate our application with very high dimensional data, although our proposed approach is sufficiently broad for general application. Specifically, the MEG data was collected from 30 HC and 30 MCI participants, using an Elekta Neuromag acquisition system with a sensor array of 102 magnetometers and 204 planar gradiometers (**Fig. 1A**, top). MEG signals were band-pass filtered (0.5-48 Hz) and segmented into nonoverlapping epochs (90 segments of 2s length). A Bayesian minimum norm was applied to estimate source time series in 8196 locations of the individual cortical surface^26^, separately for each participant (**Fig. 1A**, bottom; **Fig. 1B**, top). After the spectral analysis of the source activity using Fourier transform, FC maps were derived using the envelope of the imaginary coherence (EIC), a novel technique that we have previously developed as an alternative to traditional imaginary coherence methods^24^.

**Figure 1:**
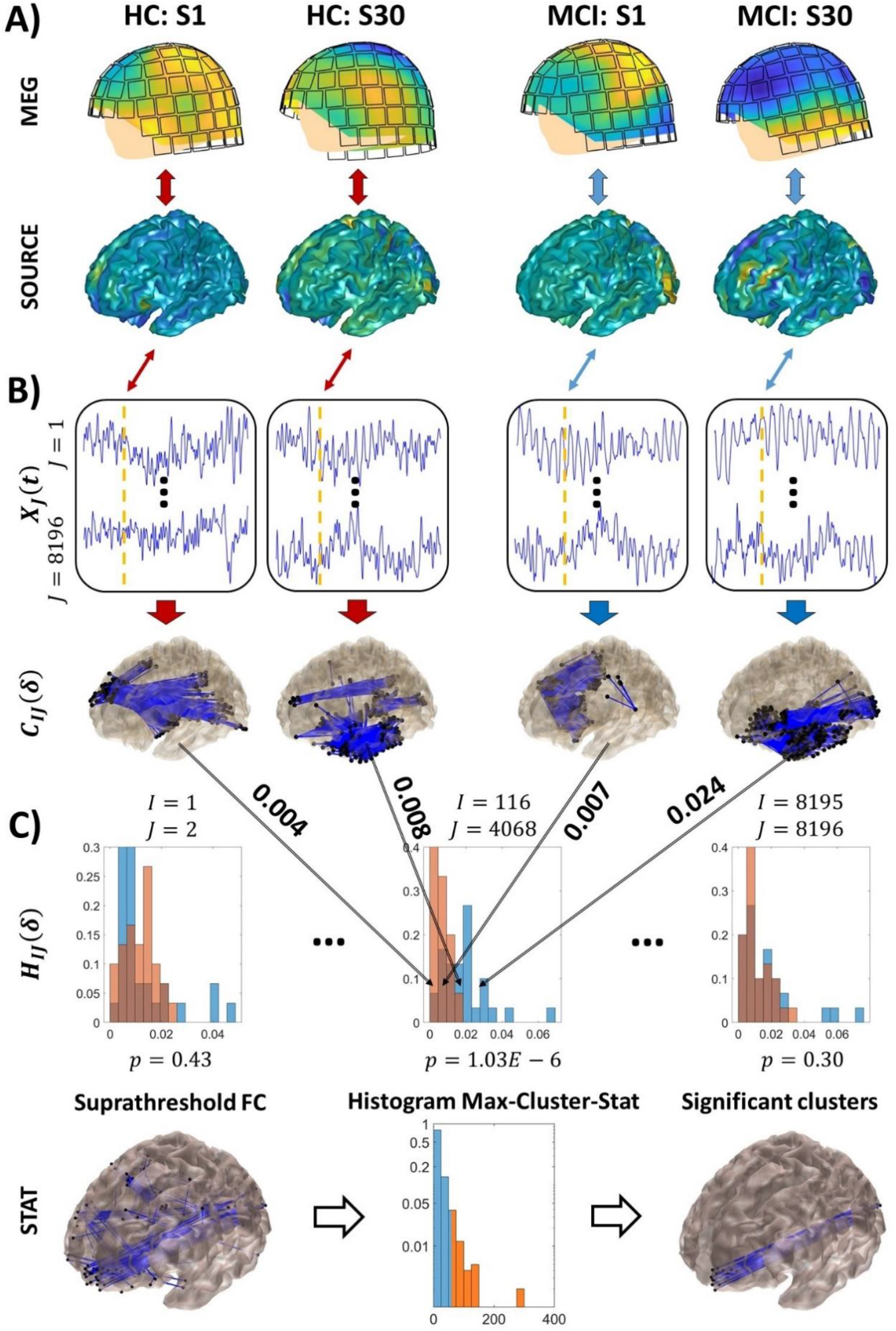
Flowchart from MEG data to functional connectivity (FC) and statistical analysis. (A) Top: MEG signals are collected from 102 magnetometers and 204 planar gradiometers, for a dataset of 30 HC (HC: S1-30, left) and 30 MCI (MCI: S1-30, right) participants. Bottom: After pre-processing, source activity is estimated using Bayesian minimum norm for source reconstruction. (B) Time-series of estimated source activity segmented into nonoverlapping epochs to produce FC maps for 8196 sources. (C) Wilcoxon rank-sum analysis of FC differences between HC and MCI participants (top). P-values are computed for these statistics and submitted to cluster-permutation statistical analysis for detecting significant clusters while controlling for multiple comparisons (bottom).

Here, the FC maps were computed directly for all the source pairs, an upper-triangular matrix of 8196 × 8195/2 elements, and averaged across frequency bands (**Fig. 1B**, bottom): 0.5-4 Hz (*δ*), 4-6 Hz (lower-theta, *θ*_1_), 6-8 Hz (upper-theta, *θ*_2_), 8-10.5 Hz (lower-alpha, *α*_1_), 10.5-13 Hz (upper-alpha, *α*_2_), 13-20 Hz (lower-beta, *β*_1_, 20-30 Hz (upper-beta, *β*_2_), and 30-48 Hz (gamma, *γ*). Thus, resulting in a total of 8 × 8196 × 8195/2 FC measures, or about 0.27 billion features. The calculations were performed separately for each participant, and the outcome consisting of a matrix of 60 rows and about 0.27 billion columns was submitted for post-hoc statistical analyses (**Fig. 1C**). Data processing steps and analyses were implemented using automated pipelines (Online Methods).

### High-dimensional FC analysis can detect brain-wide communication

The EIC method produces a normalised measure of FC strength with values between 0 and 1, similar as the coherence measure^24^. Our analyses also include neuropsychological tests scores for the assessment of participants’ cognitive abilities, namely, the Mini-Mental State Examination^27^ (MMSE), and Immediate and Delayed Recall Memory^28^ (IRM and DRM) tests. Typically, these measures have a skewed distribution, heavy tails and other features that do not comply with the normal assumption. Therefore, to avoid assumptions over the data distribution, the nonparametric Wilcoxon rank-sum test was used to compare the FC differences between HC and MCI participants, while the Spearman rank-correlation test was used to measure the monotonic relationship between the FC strength and neuropsychological tests scores.

With about 0.27 billion features, the effect size of these statistical analyses must be noticeable to be successfully measured while controlling for multiple comparisons^18^. For this purpose, we first used false discovery rate (FDR)^29^ and found significant relations in the rank-correlation analyses only between FC and DRM/IRM scores. In the analysis with DRM, we used FDR parameter *q* = 0.05 and found the significant p-values lower than 10^−7^ (*p* < 10^−7^) with correlation coefficients 0.63 < *r* < 0.69 and −0.75 < *r* < −0.63 for the positive and negative correlations, respectively. In contrast, using *q* = 0.05 in the analysis with IRM did not produce results. But we found significant results for *q* = 0.2, with *p* < 10^−6^, 0.59 < *r* < 0.69 and −0.72 < *r* < −0.59.

Above results were summarised by counting the number of significant brain-wide connections. Specifically, **Table 1** shows the outcome separately for the DRM and IRM tests, the positive and negative correlations, and for each frequency band. Notice that the number of significant negative correlations was much more prominent for lower frequencies for both cognitive tests, whereas positive correlations were more prominent for higher frequencies. Interestingly, lower values of the cognitive tests are expected for participants showing a mild or advanced stage of dementia with respect to age-matched HC. Consequently, our results showed a significant relationship between the number of significant connections, or increased FC strength, with cognitive decline, in the lower frequency bands. Thus, cortical hypersynchronization (i.e. higher FC strength in MCI with respect to HC participants) in the lower frequency bands could be associated with mild cognitive decline, as previously reported in the literature^13,30–32^.

**Table 1:** Number of significant FC links correlated with IRM and DRM scores. Positive and negative correlations are counted separately for each considered frequency band: 0.5-4 Hz (*δ*), 4-6 Hz (lower-theta, *θ*_1_), 6-8 Hz (upper-theta, *θ*_2_), 8-10.5 Hz (lower-alpha, *α*_1_), 10.5-13 Hz (upper-alpha, *α*_2_), 13-20 Hz (lower-beta, *β*_1_), 20-30 Hz (upper-beta, *β*_2_), and 30-48 Hz (gamma, *γ*). More relevant negative interactions were found in lower frequency bands, particularly *δ* and *θ*_2_ bands (highlighted in blue colour in the online version), whereas more relevant positive interactions were found in higher frequency bands, particularly *α*_2_ band (red colour).

| | FC↔DRM correlation (FDR $q = 0.05$) | | | | | | | | FC↔IRM correlation (FDR $q = 0.2$) | | | | | | | |
| --- | --- | --- | --- | --- | --- | --- | --- | --- | --- | --- | --- | --- | --- | --- | --- | --- |
| | $\delta$ | $\theta_1$ | $\theta_2$ | $\alpha_1$ | $\alpha_2$ | $\beta_1$ | $\beta_2$ | $\gamma$ | $\delta$ | $\theta_1$ | $\theta_2$ | $\alpha_1$ | $\alpha_2$ | $\beta_1$ | $\beta_2$ | $\gamma$ |
| **Positive** | 0 | 0 | 0 | 0 | 30 | 6 | 1 | 2 | 0 | 0 | 0 | 4 | 34 | 20 | 2 | 28 |
| **Negative** | 21 | 14 | 419 | 5 | 0 | 0 | 0 | 0 | 175 | 59 | 541 | 143 | 4 | 0 | 2 | 0 |

Our results are also consistent with the notion that the DRM score seems to provide a more sensitive measure of cognitive decline than IRM and other tests^33^. As mentioned above, in contrast to the analysis with DRM, no significant associations were found for the analysis with IRM when FDR was applied with *q* = 0.05. Furthermore, both DRM and IRM scores exhibited very similar trend information, as the Spearman rank-correlation analysis between both scores produced an almost perfect relationship (*r* = 0.94, with negligible p-value). To some extent, this is also consistent with our previous analysis (albeit using a different dataset) that showed a probabilistic causal relationship between immediate and delayed recall memory scores^34^.

The results for both the DRM and IRM analyses were further explored using high-dimensional FC mapping of the significantly correlated connections (**Fig. 2A**). From the FC maps, we could clearly see the increased details as compared to traditional ROI-based approaches. Our results revealed that the strongest correlations were predominantly found within a bundle of connections among occipitofrontal, occipitotemporal and parietotemporal regions in the left hemisphere in *δ* band (**Fig. 2A-B**), whereas two separated connection bundles could be observed connecting central and occipitotemporal regions in the left hemisphere in *θ*_2_ band (**Fig. 2C-D**). Significant correlations were also exhibited with connections in the right hemisphere and between both hemispheres, but these networks seemed to be much less organized in comparison with the left-hemispheric connectivity. The substantial overlap of the FC cortical maps for both the DRM and IRM corroborated the above-mentioned tight relationship between DRM and IRM scores.

**Figure 2:**
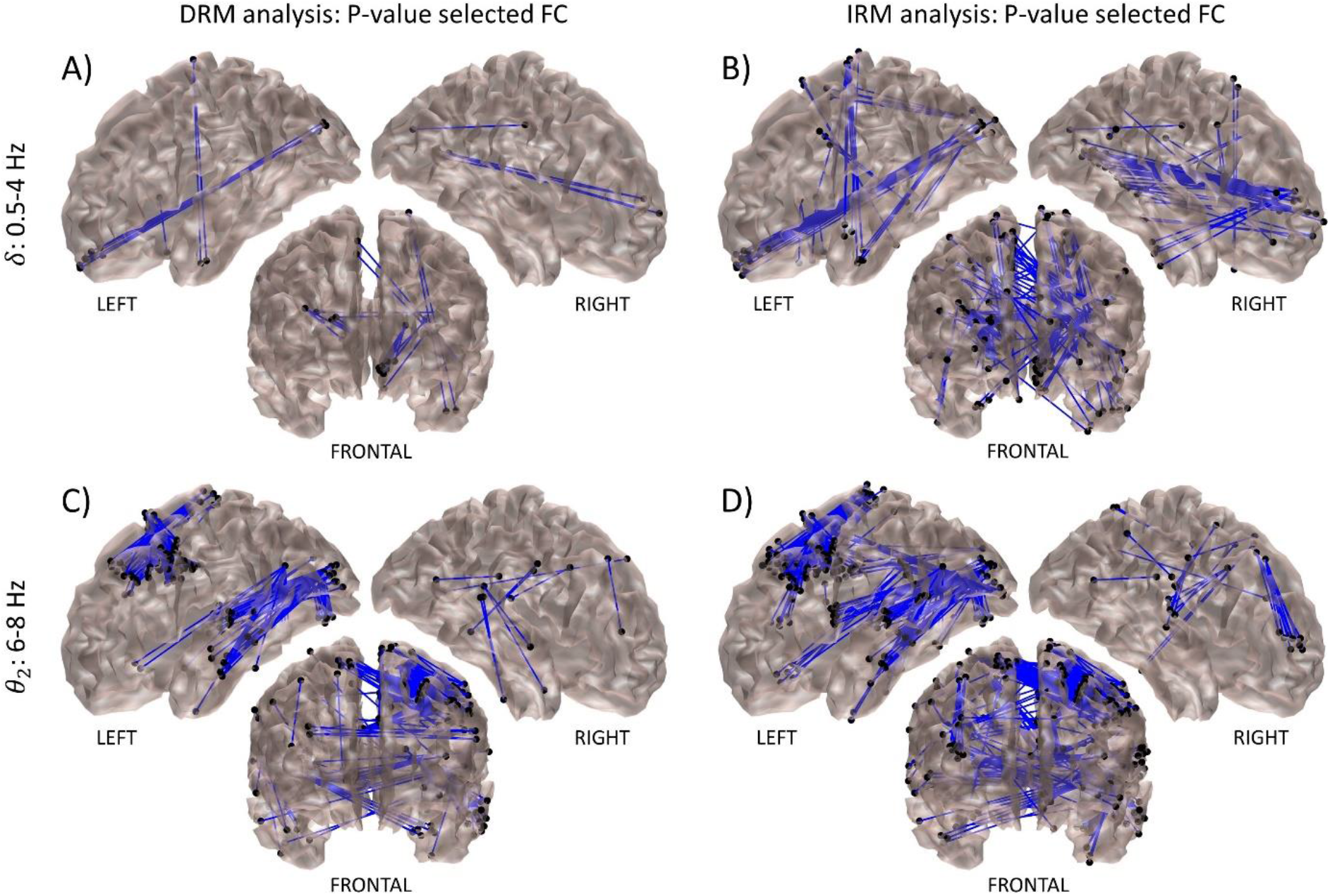
Cortical maps of FC significantly correlated with cognitive (DRM or IRM) scores. Each FC map is topographically presented in three views: left/right lateral views of the cortical hemispheres and frontal view. The left/right view only shows connections between regions in the same hemisphere, whereas the frontal view shows all the significant FC. (**A-B**) Significant FC in *δ* band (see **Table 1**). (**C-D**) Significant FC in *θ*_2_ band. Significant correlation of FC strength with cognitive scores was tested using FDR for both DRM and IRM tests, with FDR parameter *q* = 0.05 for DRM and *q* = 0.2 for IRM test. A higher value of *q* was needed for IRM test as it was less sensitive than DRM.

Next, we summarised our results using a parcellation of the cortical surface into ROIs only for comparison purposes with the literature. Specifically, we employed the Desikan-Killiany atlas^35^ and reported the significant inter-regional FC derived from the high-dimensional source FC analyses. **Figure 3** showed a schema ball summarization of the correlation analysis between the FC and DRM scores for the more relevant interactions reported in **Table 1**. In this representation, the number of connections between any two ROIs was estimated as the number of significant connections between the ROI sources. As shown in each schema, this number was normalised with respect to the highest value within each representation.

**Figure 3:**
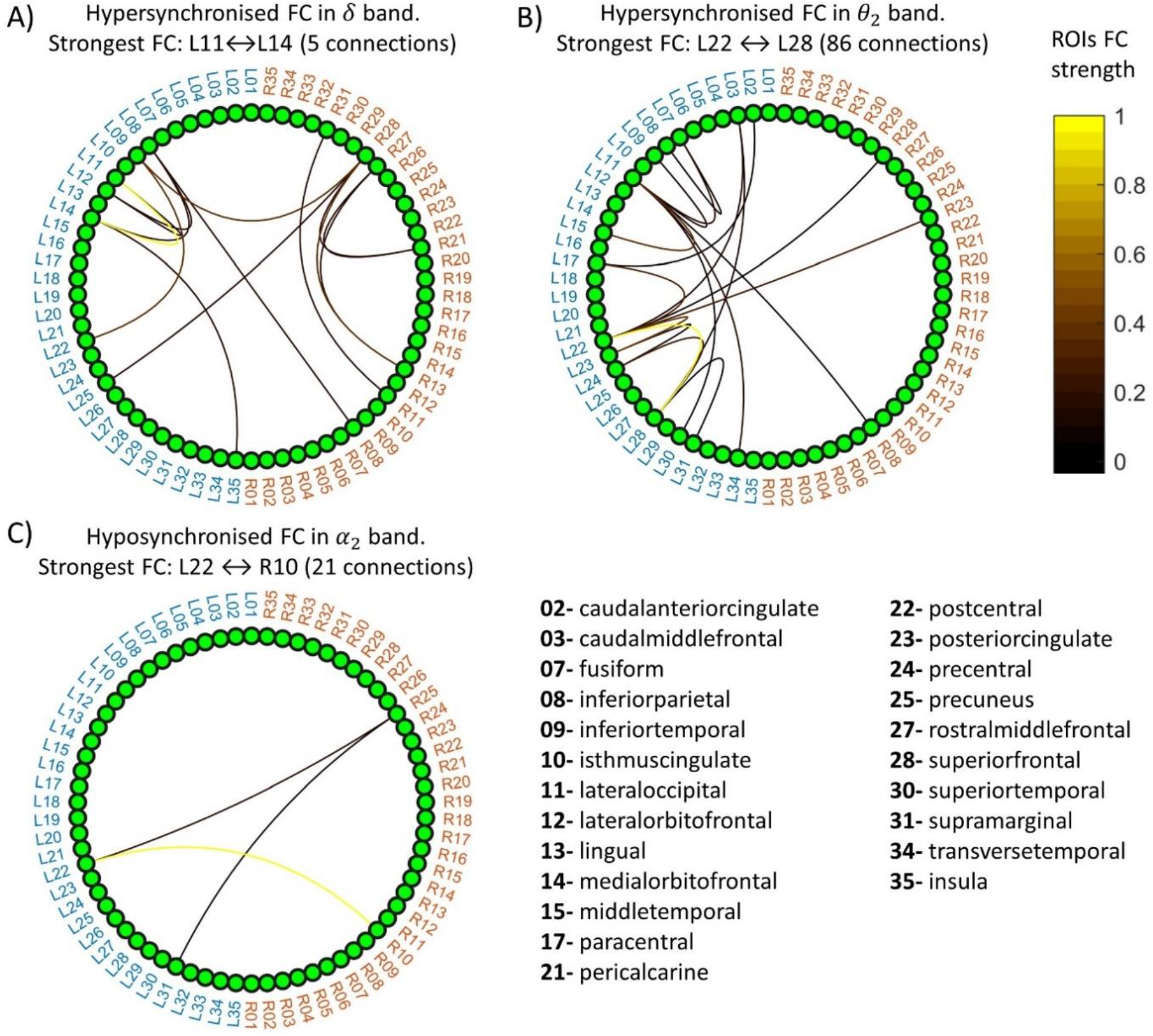
Inter-regional FC significantly correlated with DRM score. The connectivity strength between two regions of the Desikan-Killiany atlas is estimated as the number of significant connections between the two regions in the corresponding high-dimensional correlation analysis. Region labels are shown separately for the left hemisphere (blue) and right hemisphere (red). Colourmap indicates the inter-regional connectivity strength (values between 0 and 1), where this value is normalised as the number of connections divided by its highest value. (**A-B**) Graphs of hypersynchronized connectivity (FC strength significantly higher in MCI with respect to HC participants, MCI>>HC) as identified in *δ* and *θ*_2_ frequency bands. All significant connections in these bands showed negative correlations (see Table 1). (**C**) Graph of hyposynchronised connectivity (MCI<<HC) as identified in *α*_2_ frequency band. All connections in this band showed positive correlations (see Table 1).

For the FC hypersynchronization, the most prominent associations were found between the lateral occipital and medial orbitofrontal regions in *δ* band (**Fig. 3A**, 5 connections between ROI #11 and ROI #14 in the left hemisphere, or L11↔L14), and between the adjacent postcentral and superior frontal regions in *θ*_2_ band (**Fig. 3B**, L22↔L28 with 86 connections). Otherwise, the most relevant hyposynchronization (FC strength is decreased in MCI with respect to HC) was found between the left-hemispheric postcentral regions and right-hemispheric isthmus cingulate cortex in *α*_2_ band (**Fig. 3C**, L22↔R10 with 21 connections).

### Cluster-permutation statistic more consistently detects brain-wide communication

To recall, the proposed high-dimensional analyses involved about 0.27 billion features. Therefore, Bonferroni and FDR tests can be expected to produce conservative results. In this situation, it has been shown that the cluster-permutation could serve as a less conservative statistic since it can exploit the spatial structure in the data^18,19^. By clustering together spatially-related features in combination with a permutation approach that preserves the spatial structure, the cluster-permutation statistic can automatically reduce dimensionality while increasing the sensitivity of post-hoc statistical analyses.

We shall next portray its implementation by proposing a novel measure of spatial neighbourhood. Briefly, a neighbourhood relationship between any two connections, or edges, can be evaluated by requiring that the edges share one point while the other (dissimilar) endpoints are neighbours in the cortical surface (**Supplementary Fig. 2**). Then, a cluster partition of the estimated FC is created from neighbouring edges using breadth-first search (Online Methods).

The cluster-permutation statistic was computed after defining thresholds for selecting relevant features at both the lower and upper tails, separately (Online Methods). The thresholds were empirically selected as corresponding to p-values of *p*_1_ = 10^−7^, *p*_2_ = 10^−6^, and *p*_3_ = 10^−5^. This selection was based on the reasoning that with about 0.27 billion features, the expected number of spuriously selected features is about 27, 270, and 2700, correspondingly to the defined thresholds. However, the probability for these “false positive” connections to agglomerate in clusters by chance or, similarly, the probability to observe a cluster with high cardinality by chance, is expected to be much lower than for the discovered FC corresponding to the actual networks, which is the critical advantage of using cluster-permutation statistic. Moreover, we expect to obtain significant clusters of different sizes depending on the specific chosen supra-threshold value. Specifically, we could obtain narrower extended clusters for more conservative values (*p*_1_ = 10^−7^), while wider clusters could be obtained for higher values (*p*_3_ = 10^−5^).

For computing the cluster-permutation statistic using the actual data, after the pruning of edges with p-values that exceeded one of the mentioned supra-thresholds, we partitioned the surviving edges into clusters by using the defined neighbourhood measure and annotated each cluster extension (number of connections). For the identification of the significant clusters, we repeated the same procedure for randomly shuffled (surrogate) data and annotated only the maximum size of the produced clusters for each replication^18,23^. The surrogate data was generated based on the null hypothesis of having no difference between the contrast groups (MCI vs. HC) under the Wilcoxon rank-sum analysis, or no monotonic relationship between the FC strength and MMSE/DRM/IRM scores under the Spearman rank-correlation analyses. According to the hypotheses, we can assume that the labels and ranking of the measures are redundant and our data can be reshuffled^20^. As a result, we calculated a distribution for the maximum-cluster-size statistic using 1000 Monte-Carlo replicas of this process, and selected the 95^th^ percentile of this distribution as the cut-off value. Thus, this value was used to remove all the clusters with lower extension that were estimated from the actual data, thus controlling for multiple comparisons.

When we applied this technique, we obtained significant results only for the correlation analysis with DRM in the *θ*_2_ band, and for the MCI vs. HC contrast in the *δ* band. **Figure 4** showed the cortical FC maps of the whole set of connections surviving the pruning according to the defined supra-threshold p-values (**Fig. 4A**), ordered from the most (top) to less conservative (bottom) supra-thresholds. The distribution of the maximum-cluster-size is shown in the middle column together with the 95^th^ percentile of the distribution, correspondingly from top to bottom for each threshold (**Fig. 4B**). Unsurprisingly, this value increases dramatically from more conservative analysis (95^th^ percentile of maximum-cluster-size, *MCS* = 59.0) to less conservative ones (*MCS* = 603.5). As explained above, only those clusters estimated from the actual data exceeding the corresponding critical value (*MCS*) were retained (**Fig. 4C**).

**Figure 4:**
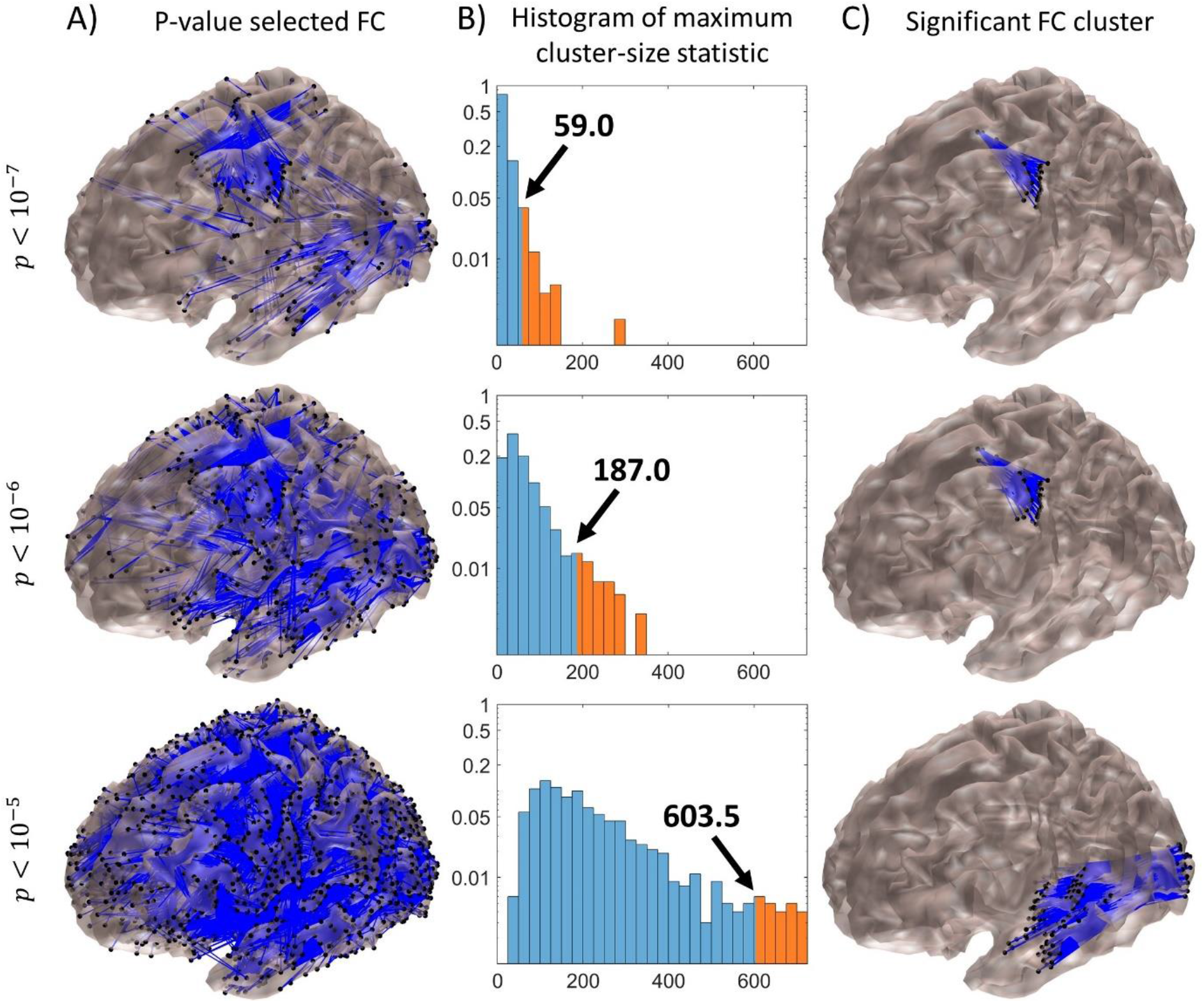
Significant clusters detected using the cluster-permutation statistic for the Spearman rank-correlation analysis between FC strength and DRM score in *θ*_2_ band. Three different supra-threshold values were tested as represented per row. (**A**) Cortical maps of all connections surviving after pruning for each supra-threshold value. (**B**) Normalised histograms of the probability distributions of the maximum-cluster-size statistic (horizontal-axis) with corresponding arrow-annotated 95^th^ percentile, which is the critical value for selecting the significant clusters in the actual data. The distribution upper tail is highlighted in orange. The vertical axis represents the relative probability values in the range 0-1, shown in a log-scale for clear visibility. (**C**) Significant clusters that remain after removing the clusters with extension lower than the corresponding critical value.

In **Figure 4C**, notice that the same cluster with different extensions, involving the communication among the central regions, was significant for the more conservative supra-threshold values (top two rows). For the most conservative (*p*_1_ = 10^−7^), it survived with extension of 135 connections (*MCS* = 59.0), whereas for a less conservative threshold (*p*_2_ = 10^−6^) it survived with extension of 227 connections (MCS = 187.0). Interestingly, for the least conservative supra-threshold (*p*_3_ = 10^−5^) this cluster vanished while a completely different cluster appeared with a much higher extension of 1019 connections (*MCS* = 603.5), which involved the communication between occipital and temporal regions in the left hemisphere (**Fig. 4C**, bottom). The results were consistent with a previous observation that more conservative thresholds could reveal spatially focal clusters, whereas the lesser conservatives reveal widely extended clusters^17,36^. Notice also that these clusters involved hypersynchronized connections indicating that an increased FC in these regions was a sign of memory impairment, as they were negatively rank-correlated with DRM (–0.75 < *r* < –0.58 for the bigger cluster of central regions, and –0.69 < *r* < –0.53 for the occipitotemporal cluster).

Complementarily, it is interesting to study the inter-group FC differences (MCI vs. HC), as it can reveal further information about the possible network changes associated with cognitive decline. Using the Wilcoxon rank-sum method within the cluster-permutation approach, we found a significant cluster of hypersynchronized FC between occipitofrontal regions in the left hemisphere in *δ* band (**Fig. 5**), i.e. communication between these regions was significantly increased in MCI with respect to HC participants. However, this cluster was only significant for *p*_2_ = 10^−6^, which further illustrates that selecting an appropriate supra-threshold value could be challenging when using the cluster-permutation technique.

**Figure 5:**
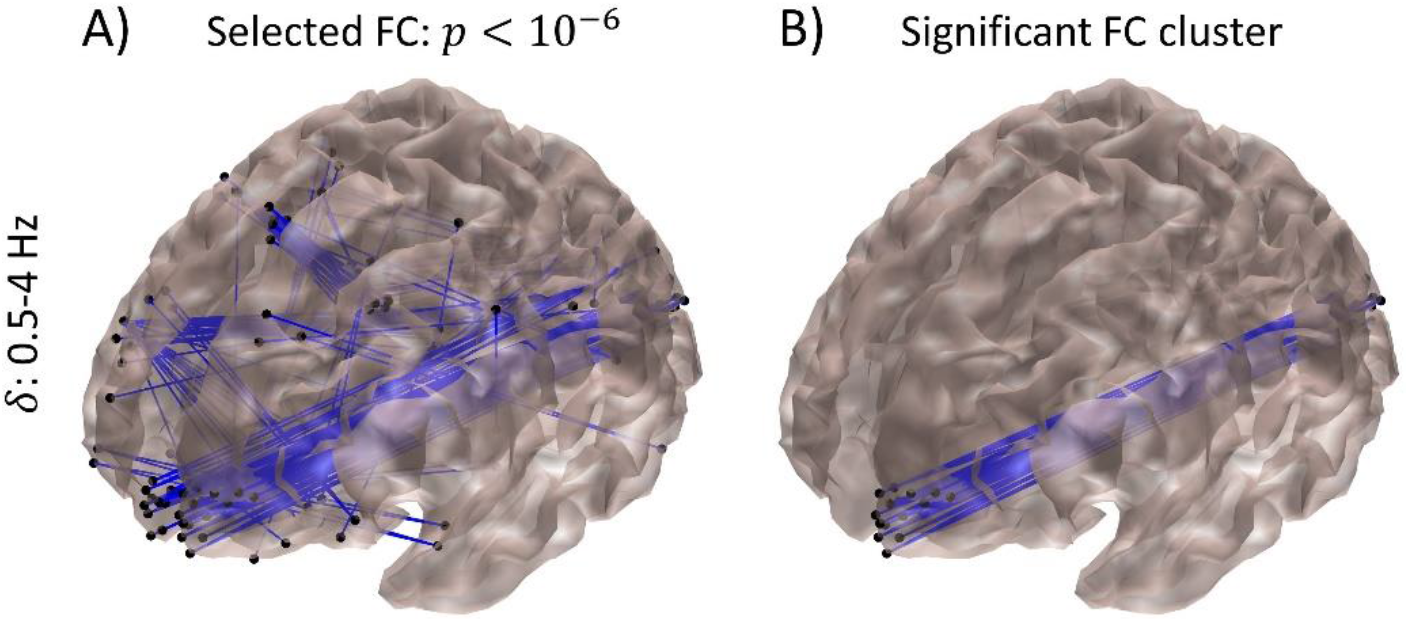
Significant clusters detected using the cluster-permutation statistic for the Wilcoxon rank-sum analysis (MCI vs. HC) in *δ* band. (**A**) Connections surviving after pruning for a selected supra-threshold value (10^−6^). (**B**) Only a cluster involving occipitotemporal regions (111 connections) survived after the correction by maximum-cluster-size statistics. See Figure 4 caption for further details.

Interestingly, in the last result, we found a nice overlap between the occipitofrontal cluster obtained with the Wilcoxon analysis (**Fig. 5B**) and the connections obtained with the Spearman analysis (**Fig. 2A**) in the same frequency band (*δ*), where the latter analysis used DRM and FDR statistic, thus demonstrating consistency. Likewise, a detailed inspection of the significant clusters obtained from the Spearman analysis with DRM also revealed consistency between the significant clusters of the central and occipitotemporal regions (**Fig. 4C**) with the significant connections obtained using the FDR statistic in *θ*_2_ band (**Fig. 2C**). Overall, this suggested the significance of these interactions at both the cluster and connection levels in our study. However, the analysis using cluster-permutation statistic has the advantage of providing a more robust evidence about the actual communication of involved brain regions.

### High-dimensional source MEG-FC analysis for neuromarkers of cognitive dysfunctions

Consistent with previous MEG-FC research on Alzheimer’s disease^32^, we consider that the hyperconnectivity patterns in the lower frequency bands could be linked to early signs of cognitive decline. In our study, we found that the hypersynchronization of critical regions in the brain, such as those involved in occipitotemporal and occipitofrontal networks, could be associated with ongoing AD pathology^37^. Thus, we shall below further investigate whether it is possible to derive a source MEG-FC neuromarker of MCI.

Beforehand, the above statistically significant clusters were mapped into ROI connectivity maps using the Desikan-Killiany atlas. Thus, the clustered brain-wide FC was used to produce a connectivity weight matrix for the atlas ROIs. **Figure 6A** exposed the weight matrix for the cluster of central regions in *θ*_2_ band, obtained from the Spearman rank-correlation analysis with DRM (**Fig. 4C**, middle row). This cluster revealed that the strongest interaction was observed between postcentral and superior frontal regions (126 connections). **Figure 6B** exposed the weight matrix for the cluster of occipitotemporal FC links in *θ*_2_ band, obtained from the same analysis (**Fig. 4C**, bottom row), where the strongest association was found between the lateral occipital area with the middle, superior, and transverse temporal regions with 340, 350 and 139 connections, respectively. **Figure 6C** exposed the weight matrix for the cluster of occipitofrontal connections in *δ* band, obtained from the Wilcoxon rank-sum analysis (**Fig. 5B**), in which the strongest FC was between the lateral occipital region with the lateral and medial orbitofrontal regions, with 25 and 69 connections, respectively. These brain regions are known to be critical for memory, emotion, object and face processing, which are among the principal cognitive functions affected during AD progression^38–42^.

**Figure 6:**
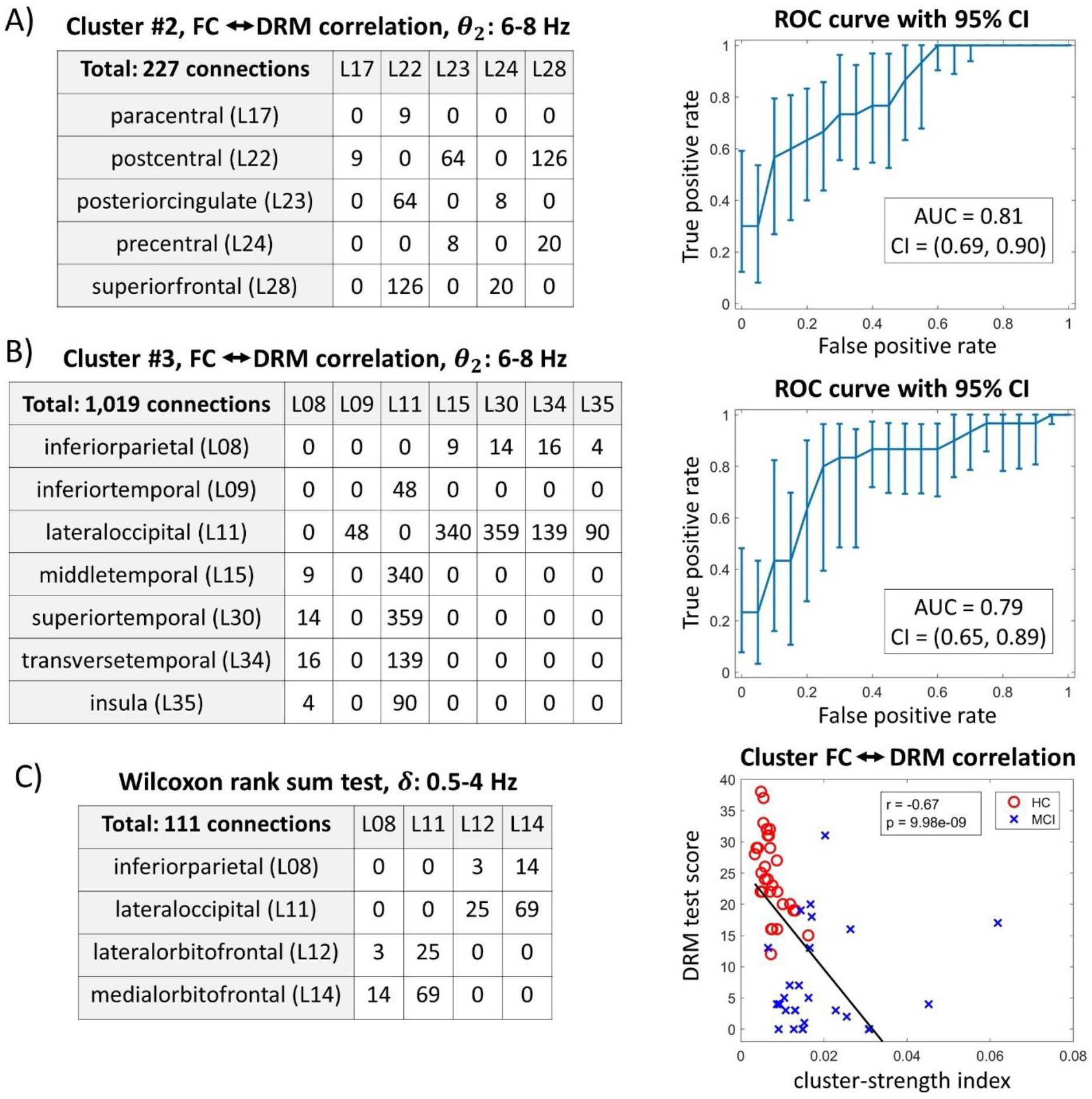
Biomarker evaluation of cluster-strength index for prediction of cognitive decline. (**A-C**) Clusters derived from Spearman rank-correlation and Wilcoxon rank-sum analyses are further scrutinised (cluster FC maps were plotted in **Fig. 4C**, middle and bottom row, and **Fig. 5B**, correspondingly in the same order). For each cluster, inter-regional weight connectivity matrices were obtained by counting the number of significant connections within two different regions of the Desikan-Killiany atlas, while a cluster-strength index was estimated as the average of all the cluster connections separately for each participant. Note that each cluster only connects a few number of ROIs as shown in the matrices. (**A**, left-side column) The first cluster mostly connects postcentral with posterior cingulate and adjacent superior frontal regions. (**B**, left-side column) The second cluster contains a hub located in the lateral occipital cortex, with a significant number of connections to inferior, middle, superior, and transverse temporal gyrus, and insula. (**C**, left-side column) The third cluster mostly connects lateral occipital with lateral and medial orbitofrontal cortex. (**A-C**, right-side column) Each cluster-strength index is used to evaluate cognitive decline. (**A-B**, right-side column) For the first two clusters, ROC analysis, with estimation of the 95% confidence interval and AUC, was conducted by providing the cluster-strength index and the corresponding HC/MCI label per participant. (**C**, right-side column) For the last cluster, Spearman rank-sum correlation analysis was conducted between the cluster-strength index and DRM test scores. Red open circle: HC participant; blue cross: MCI.

To evaluate whether these clusters could be used to predict cognitive decline, we first proceeded to average all the cluster connections to produce a single-valued cluster-strength index, separately for each cluster and participant, i.e. averaging the 227, 1019 and 111 connections, corresponding to the central (**Fig. 6A**), occipitotemporal (**Fig. 6B**) and occipitofrontal (**Fig. 6C**) networks. Subsequently, we evaluated the predictive value for the first two clusters using receiver operating characteristic (ROC) analysis. For simplicity, we provided the classification targets (HC or MCI) and the cluster-strength index as parameters for estimating the ROC curves, their 95% confidence interval (CI), and corresponding area under curve (AUC) values, using *N* = 1000 bootstrap replications. We found that the cluster-strength index showed a high classification performance with *AUC* = 0.81, *CI* = [0.69; 0.90], and *AUC* = 0.79, *CI* = [0.65; 0.89], for the corresponding clusters (**Fig. 6A-B**, right-side column). Finally, we conducted Spearman rank-correlation analysis between the third cluster-strength index and DRM test scores, which revealed a strong negative correlation with *r* = –0.67 and *p* < 10^−8^ (**Fig. 6C**, right-side column), reinforcing the view that networks hypersynchronization may be an early sign of cognitive decline. Notice that the first two clusters cannot be used in the latter analysis, nor the third cluster can be used in the former analysis, without incurring in statistical circularity. Overall, these results demonstrated the high predictive value and sensitivity of using the cluster-strength index, despite its simplicity of being just a single feature, and provided an optimistic prospect for the development of a source MEG-FC neuromarker in AD research.

## Discussion

To the best of our knowledge, this is the first time that EEG/MEG analyses have been conducted on such a large scale with about 0.27 billion features. In the current applications, FC analyses are often limited to the use of coarse brain regional parcellation, falling short of fully exploiting the ample information of the original data^1,9,16,18,32^, and thus leading to information loss, reduced statistical sensitivity, and consequently to potential bias in research findings^16,19^. To avoid such limitations, here we study the brain functional networks, with its intrinsic high-dimensional characteristics, using a robust FC measure^24^ and a novel extension of the cluster-permutation statistic^17–19^. We demonstrate our approach based on source MEG-FC analysis, using a database of 30 HC and 30 MCI participants that included MEG and neuropsychological data. The MCI participants had memory impairment and hippocampal atrophy (as detected by structural MRI scans, **Supplementary Table 1**), thereby the MCI can be linked to AD pathology with an intermediate likelihood^25^. Our application is of utmost importance to show the prospect of source MEG-FC analysis for studying neuromarkers of Alzheimer’s disease (AD) and its earlier progression.

### Extension of cluster-permutation statistical analysis for high-dimensional FC analyses

As an important contribution, we developed an extension of the cluster-permutation statistic^17–23^ to make possible our study involving a huge number of connections in source MEG-FC analysis. This was achieved by developing a new neighbourhood measure for cortico-cortical connections through the exploitation of their smooth spatial distribution patterns (see **Fig. 4,5** and Online methods). Our approach was unlike previous studies that used high-dimensional FC analysis or cluster-permutation statistic in a reduced space, by: (i) conducting the analysis among the sources of selected regions based on prior information^43^; (ii) analysing significant connected components^18^; (iii) considering the connectivity mapping of a seed point within the brain^12^; or (iv) using coarser grids to avoid the possible spurious estimation of FC among nearby sources^17,19^. Thus, we have used most of the available information in a source MEG-FC manifold to render less-biased conclusions.

Given the similarity of our approach with the spatial pairwise clustering (SPC) technique^17^, our cluster-permutation statistic could be considered as an extension of the SPC approach, especially regarding our definition of a novel cortico-cortical FC neighbourhood measure. Importantly, our work has shown that the cluster-permutation statistic can be applied to high-dimensional data without much computational burden, thus refuting previous considerations^17^ (see Online Methods). Furthermore, it should be noted that our method controls the family-wise error rate by testing significance at the cluster instead of individual connections, similar to other cluster-based permutation techniques^17–20^. Therefore, when applying our approach to source MEG-FC analyses, we focused on the significant detected FC clusters and proposed a cluster-strength index. Not only our significant clusters revealed critical networks involved in memory and cognitive processing (**Figs. 4–5**), our cluster-strength index also showed high performance and sensitivity in evaluating the cognitive status of the participants (**Fig. 6**).

### ROI vs large-scale brain-wide based FC analysis

It would have been possible to apply our approach to similar sensor-level FC analyses^32^, with the advantages that the number of sensors is considerably smaller than the number of sources. As nearby sensors record similar oscillations emanating from the underlying neuronal population, it is appropriate to use cluster statistics to exploit the smooth spatial FC patterns. However, a sensor can also identify significant activity from multiple sources located at distant sites, thus hindering the results interpretation. Less negative issues occur when using ROI-based source FC analyses. One extreme case is when the ROIs are coarsely defined by the brain lobes, in which case it resembles the sensor-level analysis. At the other extreme, FC analyses based on a very fine ROI parcellation^43,44^ could produce results comparable with those using a high-dimensional approach.

However, despite the observation that ROI analysis may provide robustness against inter-participant functional and anatomical variability^16^, it certainly leads to loss of information with the associated degraded sensitivity of post-hoc statistical analyses and biased results^16–18^. In contrast, our approach uses most of the available information in a high-dimensional manifold while dealing with volume conduction in source FC analysis^24^. Furthermore, the inter-participant variability is controlled in our approach due to the characteristic spatial blurriness of estimated source activities^26^, which may be conveyed to the FC analysis. Altogether, our cluster-permutation approach may lead to increased robustness of statistical analyses, and the significantly detected FC clusters should reasonably increase our confidence in having discovered actual brain networks.

In contrast to ROI analysis, the use of large-scale brain-wide FC approach also allows for direct comparisons with ECoG analysis to measure FC in the brain^12,45^. Particularly, with the use of the EIC method or an imaginary coherence index^24,46^, which are robust to volume conduction, we ameliorate the risk that the measured interactions are spurious and allow for the robust estimation of short-range connectivity in the brain. Critically, EIC can also measure true interactions caused by zero-lag (modulus *π*) phase interactions that are neglected by other imaginary coherence measures^24^, which is an advantage inherited by our implementation.

### Prospective neuromarker importance of source MEG-FC analysis

Consistent with our findings (**Figs. 2–6** and **Table 1**), the state-of-the-art in AD research using EEG/MEG data have established a relationship between hypersynchronized FC and memory decline^10,13,14,31,32^. Particularly, our high-dimensional FC analyses exposed significant hypersynchronized communication of occipitotemporal and occipitofrontal regions (**Fig. 6**, left-side column). Such hypersynchronization phenomenon could be attributed to overall neuronal excitatory enhancement^47^, reduced disinhibition of neurons^30^, or as a compensatory mechanism related to brain plasticity changes triggered by AD synaptic and neuronal loss^48,49^.

Among the significantly found connected regions, there is a consensus that the insula and temporal areas (e.g. transverse temporal, **Fig. 6B**) are critically involved in episodic memory processes^38,39^. Other significantly connected regions in the frontal (lateral and medial orbitofrontal cortex), lateral occipital, and inferior parietal cortices (**Fig. 6C**), are important for decision making, reward evaluation, face/object and emotion processing^40–42^.

Furthermore, our main results (**Figs. 4–6**) overlapped with previously reported hypersynchronization in similar brain regions^14,15,32^. Although, in our case, the identification of significant FC clusters provides stronger evidence for this claim, as clusters can more robustly support the evidence of inter-regional communication in contrast to the sparse FC patterns exposed in previous works^13–16,32^, with the additional advantage that the smooth characteristic of FC clusters also brings robustness against inter-subject anatomical and functional variability. Therefore, we suggest that these two properties of FC clusters are important for a source MEG-FC neuromarker in dementia. Here, we further supported the importance of cluster analysis by showing that a cluster-strength index can be used to evaluate cognitive decline with promising results (**Fig. 6**, right-side column). However, note that these results must be interpreted with caution because the limited sample size in our study. Thus, our results must be further scrutinised using different databases in future studies.

Overall, we have successfully developed a novel analytical pipeline involving cluster-permutation statistics for analysing high-dimensional brain-wide FC maps, while avoiding the biases that come along with standard brain parcellation approaches. Such high-resolution FC maps can be estimated without high computational cost (see Online Methods) and could be important to advance research in healthy and unhealthy neural information processing. Our proposed approach can, in general, be applied to a variety of neuroimaging studies, including translational clinical research.

## Data Availability

The data of the present study would be available through an institutional repository and under a previous request to the authors.

## Code Availability

The MATLAB code is available at the following GitHub repository: https://github.com/JMSBornot/High-Dimensional-Source-MEG-FC

## Acknowledgments

This work was supported by the EU’s INTERREG VA Programme, managed by the Special EU Programmes Body (SEUPB) (J.M.S.-B., P. M. and K.W.-L.), the Northern Ireland Functional Brain Mapping Project (1303/101154803) funded by Invest NI and Ulster University (S.Y., G.P. and K.W.-L.), the Spanish Ministry of Economy and Competitiveness (PSI2009-14415-C03-01) and Madrid Neurocenter (M.E.L., R.B. and F.M.), Alzheimer’s Research UK (ARUK) Pump Priming Awards (G.P., P. M. and K.W.-L.), and Medical College of Wisconsin (V.Y.). P. M. and K.W.-L. received additional support from Ulster University Research Challenge Fund, and Global Challenges Research Fund, and K.W.-L. from COST Action Open Multiscale Systems Medicine (OpenMultiMed) supported by COST (European Cooperation in Science and Technology). The views and opinions expressed in this paper do not necessarily reflect those of the European Commission or the Special EU Programmes Body (SEUPB).

## Contributions

J.M.S-B. and K.W-L. formulated the idea. M.E.L., R.B. and F.M. collected the data. R.B and V.Y suggested the use of cluster statistic. J.M.S-B. devised the extension of the cluster-permutation approach. J.M.S-B implemented the methods and experiments. M.E.L., R.B., V.Y., S.Y., P.M., F.M. and G.P. provided additional suggestions relevant to the experiments and analysis. J.M.S-B. and K.W.-L. analysed the data and prepared the original draft of the paper. All authors discussed the results, reviewed and edited the paper.

## Competing interests

The authors declare no competing financial interests.

## Online methods

### Participants

Data were collected from a total of 60 participants at Hospital Universitario de San Carlos (Madrid, Spain), including MEG recordings and neuropsychological tests scores to evaluate cognitive and memory abilities (MMSE Spanish version^27^, and DRM/IRM scores from Wechsler Memory Scale-III^28^). Inclusion criteria: recruitment age of 65-85 years, right-handed as verified using Edinburgh Handedness Inventory^50^, native Spanish speakers, a modified Hachinski score ≤ 4^51^, a Geriatric Depression Scale short-form score ≤ 5^52^, and no indication of comorbidities or brain trauma according to MRI inspection^31^. MCI participants showed signs of hippocampal atrophy as quantified using their anatomical MRI, and therefore it was considered that their cognitive impairment was related to AD pathology with an intermediate likelihood^25^. HC group: N=30, 16 females, ages 66-80 years. MCI group: N=30, 15 females, ages range 65-78 years (see **Supplementary Table 1** for further details).

### MEG data recording

The MEG signals were acquired using an Elekta-Neuromag system with 306 channels (102 magnetometers and 204 planar gradiometers) with a sampling frequency of 1 kHz and online anti-alias filter with 0.1-330 Hz bandwidth. Concurrently, head movements were tracked using a continuous head-position indicator (cHPI) with four coils attached to the scalp. The position for these coils, fiducial points (nasion, and left/right preauricular), and head-shape model were digitised using a three-dimensional Fastrak Polhemus system (Polhemus, Inc, USA). Additionally, bipolar electro-oculogram sensors were attached above and below the left eye to measure ocular movements, and an electrical ground electrode was attached to the earlobe. Offline, MEG signals were processed using the temporal extension of the signal-space separation technique (Maxfilter version 2.2, Elekta; correlation threshold = 0.9, time window = 10 s) to reduce the contribution of external magnetic field and correct the head movements using the cHPI data. About 15 minutes were recorded for each participant in eyes-closed resting-state condition, although only a continuous segment of 180-seconds, relatively free of artefacts, was selected by visual inspection for post-hoc analyses.

### Pipeline for pre-processing

Analyses were conducted using MATLAB custom code. First, the individual anatomical MRI was co-registered with corresponding MEG fiducials and head-shape model points using a modified interface of the SPM12 automatic routine to improve co-registration accuracy (see **Supplementary Fig. 4**). Subsequently, the MEG lead field was estimated using the SPM12 implementation of the single-shell technique with sources located on the SPM12 template surface of 8198 vertices (medium size). Similarly, using the SPM12 toolbox within a MATLAB custom script, signals were pre-processed using a Butterworth’s bandpass filter of 0.5-48 Hz bandwidth, downsampled to 200 Hz and epoched into 2-seconds segments (90 trials) using a Hann window. Next, the source reconstruction analysis was conducted using the SPM12 implementation of the Bayesian minimum norm optimization^26^ for each segment. Finally, the discrete Fourier transform (MATLAB fft function) was applied to each estimated and segmented source activity, and its derived complex numbers were halved to “single” precision and saved to hard-disk for post-hoc FC analyses. In summary, this resulted in a 3D matrix of dimensions 96 frequency bins (frequency resolution of 0.5 from 0.5 to 48 Hz), 8196 sources and 90 segments, for each subject.

### Pipeline for FC analysis

Specifically, in this study the envelope of the imaginary coherence (EIC)^24^ was estimated as the FC index between two signals *x_i_*(*t*) and *x_j_*(*t*), for each frequency *ω*:

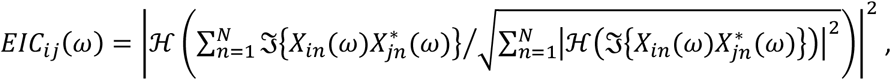

where *X_in_*(*ω*) is the Fourier transform complex number output for signal *x_in_*(*t*), estimated separately for each epoch *n* = 1, …, 90; the operator 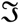 extracts the imaginary part of the argument’s complex number; 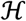 represents Hilbert’s transform; and |·| is the absolute value.

Due to limited RAM memory for conducting the statistical analyses, it is troublesome to compute the whole connectivity matrix of 8196×8196 interactions. Thus, we partitioned the FC matrix into a 16×16 sub-block matrices for a total of 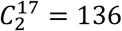 blocks (15 blocks times 500 sources + 1 block times 696 sources = 8196). We only kept the block subindices for the strictly upper triangular part due to the symmetry of the FC measure (**Supplementary Fig. 1**). After the block-wise FC estimation, the EIC values were averaged for the interested frequency bands. These bands were chosen by modifying the classical EEG/MEG band partitioning to allow a slightly higher level of detail (delta *δ*: 0.5-4 Hz, lower-theta *θ*_1_: 4-6 Hz, upper-theta *θ*_2_: 6-8 Hz, lower-alpha *α*_1_: 8-10.5 Hz, upper-alpha *α*_2_: 10.5-13 Hz, lower-beta *β*_1_: 13-20 Hz, upper-beta *β*_2_: 20-30 Hz, and gamma *γ*: 30-48 Hz). Finally, the FC estimates were saved to hard-disk, separately for each participant, frequency band and block, for post-hoc statistical analyses.

### Non-parametric statistical analyses

Our study involves a matrix data for 60 participants (rows) and about 0.27 billion features (columns), where the features corresponded to the estimation of 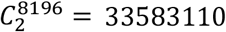 pairwise FC for each of the above-mentioned frequency bands, together with about 60 cognitive tests scores collected from the participants during MMSE, IRM and DRM tests (one or two missing data in each test). These measurements are used for: (i) the Wilcoxon rank-sum analysis of the differences between 30 HC and 30 MCI participants for each feature; and (ii) the Spearman rank-correlation analyses between each cognitive test and feature.

In our case, the estimated FC and cognitive scores are nonnegative and hardly follow the normality assumption. Thus, we adopted a whole non-parametric approach to avoid assumptions on data distribution, and to implement the cluster-permutation approach. Non-parametric tests not only can produce more accurate results than comparable traditional techniques, e.g. Wilcoxon and Spearman tests^53^, but also are often used to exploit relevant data structure, such as when using permutation techniques to exploit spatial smoothness^17–20^.

The permutation technique is adopted here to create surrogate data under the null-hypotheses of no group differences or no correlation between the ranks of each feature and cognitive scores^20^, which is a critical step for testing the significance of FC clusters as discussed in the next section. Simply, each surrogate data is created by randomly reshuffling the row-order of the above matrix in Wilcoxon analysis, or the order of the cognitive measurements in Spearman analysis (*N* = 1000 Monte Carlo simulations). Notice that all the elements in each row of the matrix are jointly reshuffled to avoid destroying the data structure in the Wilcoxon analysis. Particularly, the row reshuffling corresponds to randomly assigning each subject to either the HC or MCI group for the creation of each surrogate data, accordingly to the hypothesis of no group differences^20^.

The implementation of our approach is a computational challenge because the Wilcoxon and Spearman analyses produce an array of 0.27 billion p-values for the original and each of the 1000 surrogate data. Therefore, we adopted the supra-threshold technique^17–20^ to select only those features with corresponding p-values lower or equal than a threshold of *p*_1_ = 10^−7^, *p*_2_ = 10^−6^, or *p*_3_ = 10^−5^. Then, for 0.27 billion features, our expected number of false alarms is 27, 270, or 2700, for each threshold. Subsequently, supra-threshold features were mapped into supra-threshold FC, separately for each frequency band, with supra-threshold FC indices in the range 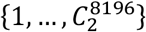, and the sparse arrays of supra-threshold connection indices were saved to hard-disk for the posterior cluster-permutation analysis.

Furthermore, recall that above analyses were conducted for the upper triangular part of the 8196×8196 FC matrix and, particularly, after partitioning it into 136 sub-block matrices, the statistical analyses were conducted block-wise due to RAM limitations. Afterwards, we loaded the sparse array of supra-threshold FC indices for each block, separately for each frequency band, and assembled the indices for all the blocks before running the cluster parcellation procedure, which will be discussed in the next section. As a summary, this first stage of the implementation of our approach can be presented as follows:

1. FC data, or features, were loaded for all the participants, separately for each block and frequency band;
2. Wilcoxon and Spearman analyses, which were based on each of the 0.27 billion features and cognitive scores, were conducted for the original and each of the 1000 surrogate data, thus producing the corresponding p-values for each feature.
3. The supra-threshold values *p*_1_ = 10^−7^, *p*_2_ = 10^−6^, or *p*_3_ = 10^−5^, were used separately for pruning off the less relevant features.
4. Only the sparse array of indices corresponding to the supra-threshold features, were saved to hard-disk for the posterior cluster statistical analysis.

### Cluster-permutation statistical analysis

We have proposed an extension and improvement of the original spatial pairwise clustering (SPC) technique^17^. Particularly, we defined a novel cortico-cortical FC neighbourhood measure for parcelling connections into FC clusters in the cortical surface, as will be explained below in more detail. The last stage of our statistical analysis can be implemented, following from the last step of the above-discussed first stage, as follows:

1. Separately for each frequency band and each supra-threshold value *p*_1_ = 10^−7^, *p*_2_ = 10^−6^, or *p*_3_ = 10^−5^, we loaded the sparse array of supra-threshold FC indices for each block, and assembled the indices for all the blocks.
2. Using the assembled FC indices, FC clusters were estimated using a breadth-first search algorithm (see **Supplementary Table 2** for a practical implementation) for the original and 1000 surrogate data.
3. The number of connections, or cluster size, was computed for each cluster of the original and surrogate data.
4. The maximum-cluster-size statistic was estimated as the maximum size among all the clusters that were estimated for each surrogate data, thus rendering a distribution of 1000 samples of this statistic in our analysis.
5. The 95^th^ percentile of the maximum-cluster-size distribution was selected as a critical value.
6. Finally, using the cluster-permutation statistic, the significant FC clusters in the original data are those clusters with size greater or equal than the critical value.

In the above implementation of our cluster-permutation approach, when we use the supra-threshold technique, not only do we apply it separately for each frequency band, but we also test independently the significance of clusters for positive and negative effects, i.e. for statistic with p-values in the upper and lower tails of the statistical distribution, in contrast to previous applications that proposed using the supra-threshold technique on the statistic absolute value^23,43^. Notice that the latter choice implies the union of clusters from measured positive and negative effects, and therefore may have a negative impact on the results. As a practical solution, the maximum-cluster-size distribution was estimated and the 95^th^ percentile of each distribution was selected, separately for each case, resulting in a set of critical values. Finally, the maximum critical value is selected in our implementation of the cluster-permutation statistic to effectively detects significant FC clusters while controlling for multiple comparison^17–20^.

### Measure of cortico-cortical FC neighbourhood

Next, we present a novel measure of neighbourhood between a pair of connections (*X*_*I*_1_(*k*)_, *X*_*I*_2_(*k*)_), *I*_1_(*k*) < *I*_2_(*k*) and (*X*_*I*_1_(*l*)_, *X*_*I*_2_(*l*)_), *I*_1_(*l*) < *I*_2_(*l*), which are defined correspondingly for the strictly upper-part of a triangular matrix that is representing a symmetric FC measure, where the unique connections are arranged using an array of FC indices 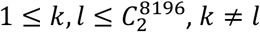, and 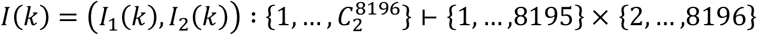 is a functional mapping of the connection to its vertices index. Thus, 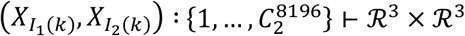.

Based on this definition, our FC neighbourhood measure can be represented as follows:

1. Check whether *I_m_*(*k*) is equal to *I_n_*(*l*) for some *m*, *n* ∈ {1,2}.
2. If true, then the two connections, or edges, have a vertex in common. Set *J*(*k*) and *J*(*l*) as the complementary vertices in the edges (*J*(*k*) = *I*_3–*m*_(*k*) and *J*(*l*) = *I*_3–*n*_(*l*)). Notice that *J*(*k*) ≠ *J*(*l*) by definition.
3. The connections are neighbours if the vertices *X*_*J*(*k*)_ and *X*_*J*(*l*)_ are neighbours in the cortical surface (**Supplementary Fig. 2**).

### Efficient computation of statistics within the permutation approach

In the implementation of our cluster-permutation statistic, we use some tricks for dealing efficiently with the calculations within the permutation procedure. For example, when running the Wilcoxon rank-sum or Spearman rank-correlation analysis, the statistic ranks should have to be computed inside the whole array of *M* = 60 participant measurements, separately for each permutation (naively). However, that will be inefficient as it involves an order of *O*(*M* log(*M*) *N*) operations. Because measurements are fixed along different permutations, where the only change is in their ranking order that change with each new permutation, then the sorted rank order could be estimated only once for the sorted statistic at the initial step, and the ultimate ranking values could be updated for each permutation, with a lower cost of *O*(*MN*) operations for all the *N* permutations (see **Supplementary Table 3** for a MATLAB code with the implementation of this idea). Similarly, our Spearman rank-correlation analysis is based on the measurements ranks, thereby the rank estimation could be optimised as previously.

Furthermore, all the computations involved in the correlation formula do not need to be undertaken for each permutation. Clearly, as this formula can be expressed as

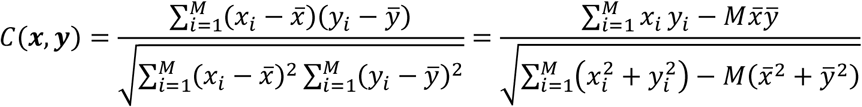

then the numerator term 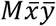, and the whole denominator can be computed once. The only term that needs to be recomputed for each permutation is 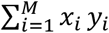. In summary, the intertwining of statistics and permutation calculations is feasible and has a significant impact on the speed of the whole procedure.

### Computational tricks for implementing cluster-permutation statistical analysis

We refuted a pessimistic observation in the SPC paper^17^, where the authors stated that the cluster partition of supra-threshold connections is “performed by initializing an *N*(*N* – 1)/2 × *N*(*N* – 1)/2 adjacency matrix”, which will be unfeasible in our case with *N* = 8196 sources. As has been shown here (e.g. see pseudocode in **Supplementary Table 2**), our testing of a neighbourhood relationship between two connections exclusively rely on the testing of a neighbourhood relationship in the cortical surface (**Supplementary Fig. 2**). Therefore, if we use an adjacency matrix to reflect this relation, our matrix will be of dimensions *N* × *N*, which is a huge improvement with respect to the original implementation. However, we recommend using a list of neighbour vertices for each vertex to test this relationship, with higher computational memory efficiency.

Furthermore, as was mentioned above, the permutation and statistical analyses were performed separately for each of the 136 sub-blocks of the 8198×8196 FC matrix and defined frequency bands. If users have a lower RAM memory capacity, a finer sub-blocks partition could be considered, i.e. a 41×41 partition for a total of 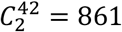 blocks (40 blocks times 200 sources + 1 block times 196 sources = 8196). Using the above supra-threshold values, only the indices of these FC measures that exceeded the corresponding critical value are saved to hard-disk, for each block and for all the permutations simultaneously. Notice that the selected indices have a sparse structure as indices for corresponding p-values exceeding either *p*_1_ = 10^−7^, *p*_2_ = 10^−6^, or *p*_3_ = 10^−5^, separately, were removed. Therefore, the expected sparse density of saved indices is 10^−7^, 10^−6^, and 10^−5^, respectively. Despite having about 0.27 billion features and running the cluster-permutation procedure for the original and 1000 surrogate data, this operation is feasible because of the mentioned high sparsity.

Finally, another apparently less significant but very important trick is to use the corresponding statistic supra-threshold values, instead of the supra-threshold p-values. Therefore, during the implementation of the cluster-permutation procedure, we avoid estimating p-values for the involved statistical analysis. Interestingly, the most important aspect of this trick is that usually p-values for the Wilcoxon and Spearman analyses are obtained using approximations because of the computational cost of using exact p-value computation. In our work, we created a lookup table for the Wilcoxon rank-sum analysis, which allowed to obtain the needed supra-threshold statistic values. For example, for our rank-sum statistic involving 30 HC vs. 30 MCI, we used the table values of 579, 603 and 629 for the lower tail (considering only the negative median differences), and 1251, 1227 and 1201 for the upper tail (considering only the positive median differences), correspondingly to the supra-threshold p-values *p*_1_ = 10^−7^, *p*_2_ = 10^−6^, or *p*_3_ = 10^−5^, separately (**Supplementary Fig. 3**). For the Spearman rank-correlation analysis, due to the high number of participants (*M* = 60), it is difficult to obtain an exact p-values lookup table as the exact method involves an order of factorial of *M* operations. Therefore, for simplicity, for the Spearman analysis we used supra-threshold correlation values that were estimated for the corresponding supra-threshold p-values, using the standard p-value approximation for the Pearson’s correlation coefficient that is based on the Student’s t-distribution, i.e. 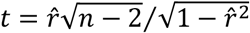, where 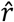 is the estimated correlation coefficient, and the p-value is estimated as 2*P*(*T* > *t*), where *T* follows a t-distribution with *n* – 2 degrees of freedom.

## References

1. van den Heuvel, M. P. & Hulshoff Pol, H. E. Exploring the brain network: A review on resting-state fMRI functional connectivity. Eur. Neuropsychopharmacol. 20, 519–534 (2010).

2. Greicius, M. D., Srivastava, G., Reiss, A. L. & Menon, V. Default-mode network activity distinguishes Alzheimer’s disease from healthy aging: Evidence from functional MRI. Proc. Natl. Acad. Sci. 101, 4637–4642 (2004).

3. de Vos, F. et al. A comprehensive analysis of resting state fMRI measures to classify individual patients with Alzheimer’s disease. Neuroimage 167, 62–72 (2018).

4. Greicius, M. D., Krasnow, B., Reiss, A. L. & Menon, V. Functional connectivity in the resting brain: a network analysis of the default mode hypothesis. Proc. Natl. Acad. Sci. U. S. A. 100, 253–8 (2003).

5. Buckner, R. L. et al. Cortical Hubs Revealed by Intrinsic Functional Connectivity: Mapping, Assessment of Stability, and Relation to Alzheimer’s Disease. J. Neurosci. 29, 1860–1873 (2009).

6. Haak, K. V., Marquand, A. F. & Beckmann, C. F. Connectopic mapping with resting-state fMRI. Neuroimage 170, 83–94 (2018).

7. Buckner, R. L., Krienen, F. M. & Yeo, B. T. T. Opportunities and limitations of intrinsic functional connectivity MRI. Nat. Neurosci. 16, 832–837 (2013).

8. Power, J. D. et al. Functional network organization of the human brain. Neuron 72, 665–78 (2011).

9. Raichle, M. E. The Brain’s Default Mode Network. Annu. Rev. Neurosci. 38, 433–447 (2015).

10. Maestú, F. et al. A multicenter study of the early detection of synaptic dysfunction in Mild Cognitive Impairment using Magnetoencephalography-derived functional connectivity. NeuroImage Clin. 9, 103–109 (2015).

11. Logothetis, N. K. What we can do and what we cannot do with fMRI. Nature 453, 869–878 (2008).

12. Hipp, J. F., Hawellek, D. J., Corbetta, M., Siegel, M. & Engel, A. K. Large-scale cortical correlation structure of spontaneous oscillatory activity. Nat. Neurosci. 15, 884–890 (2012).

13. Koelewijn, L. et al. Oscillatory hyperactivity and hyperconnectivity in young APOE-ε4 carriers and hypoconnectivity in Alzheimer’s disease. Elife 8, 1–25 (2019).

14. Dimitriadis, S. I. et al. How to build a functional connectomic biomarker for mild cognitive impairment from source reconstructed MEG Resting-state activity: The combination of ROI representation and connectivity estimator matters. Front. Neurosci. 12, 1–21 (2018).

15. Yu, M. et al. Selective impairment of hippocampus and posterior hub areas in Alzheimer’s disease: An MEG-based multiplex network study. Brain 140, 1466–1485 (2017).

16. Hillebrand, A., Barnes, G. R., Bosboom, J. L., Berendse, H. W. & Stam, C. J. Frequency-dependent functional connectivity within resting-state networks: An atlas-based MEG beamformer solution. Neuroimage 59, 3909–3921 (2012).

17. Zalesky, A., Cocchi, L., Fornito, A., Murray, M. M. & Bullmore, E. Connectivity differences in brain networks. Neuroimage 60, 1055–1062 (2012).

18. Zalesky, A., Fornito, A. & Bullmore, E. T. Network-based statistic: Identifying differences in brain networks. Neuroimage 53, 1197–1207 (2010).

19. Zalesky, A., Fornito, A., Egan, G. F., Pantelis, C. & Bullmore, E. T. The relationship between regional and inter-regional functional connectivity deficits in schizophrenia. Hum. Brain Mapp. 33, 2535–2549 (2012).

20. Hayasaka, S. & Nichols, T. E. Validating cluster size inference: random field and permutation methods. Neuroimage 20, 2343–2356 (2003).

21. Smith, S. M. & Nichols, T. E. Threshold-free cluster enhancement: Addressing problems of smoothing, threshold dependence and localisation in cluster inference. Neuroimage 44, 83–98 (2009).

22. Zhang, F. et al. Suprathreshold fiber cluster statistics: Leveraging white matter geometry to enhance tractography statistical analysis. Neuroimage 171, 341–354 (2018).

23. Maris, E. & Oostenveld, R. Nonparametric statistical testing of EEG-and MEG-data. J. Neurosci. Methods 164, 177–190 (2007).

24. Sanchez Bornot, J. M., Wong-Lin, K. F., Ahmad, A. L. & Prasad, G. Robust EEG/MEG Based Functional Connectivity with the Envelope of the Imaginary Coherence: Sensor Space Analysis. Brain Topogr. 0, 1–22 (2018).

25. Albert, M. S. et al. The diagnosis of mild cognitive impairment due to Alzheimer’s disease: Recommendations from the National Institute on Aging-Alzheimer’s Association workgroups on diagnostic guidelines for Alzheimer’s disease. Alzheimer’s Dement. 7, 270–279 (2011).

26. Mattout, J., Phillips, C., Penny, W. D., Rugg, M. D. & Friston, K. J. MEG source localization under multiple constraints: An extended Bayesian framework. Neuroimage 30, 753–767 (2006).

27. Lobo, A., Escobar, V., Ezquerra, J. & Seva Díaz, A. El Mini-Examen Cognoscitivo”(Un test sencillo, práctico, para detectar alteraciones intelectuales en pacientes psiquiátricos). Rev. Psiquiatr. y Psicol. Médica (1980).

28. Wechsler, D. Wechsler memory scale-III Manual. San Antonio, TX Psychol. Corp. (1997).

29. Genovese, C. R., Lazar, N. A. & Nichols, T. Thresholding of statistical maps in functional neuroimaging using the false discovery rate. Neuroimage 15, 870–878 (2002).

30. Garcia-Marin, V. et al. Diminished perisomatic GABAergic terminals on cortical neurons adjacent to amyloid plaques. Front. Neuroanat. 3, 28 (2009).

31. López, M. E. et al. Alpha-band hypersynchronization in progressive mild cognitive impairment: a magnetoencephalography study. J. Neurosci. 34, 14551–14559 (2014).

32. Engels, M. M. A. et al. Alzheimer’s disease: The state of the art in resting-state magnetoencephalography. Clin. Neurophysiol. 128, 1426–1437 (2017).

33. Welsh, K., Butters, N., Hughes, J., Mohs, R. & Heyman, A. Detection of abnormal memory decline in mild cases of Alzheimer’s disease using CERAD neuropsychological measures. Arch. Neurol. 48, 278–281 (1991).

34. Ding, X. et al. A hybrid computational approach for efficient Alzheimer’s disease classification based on heterogeneous data. Sci. Rep. 8, 9774 (2018).

35. Desikan, R. S. et al. An automated labeling system for subdividing the human cerebral cortex on MRI scans into gyral based regions of interest. Neuroimage 31, 968–980 (2006).

36. Friston, K. J., Worsley, K. J., Frackowiak, R. S. J., Mazziotta, J. C. & Evans, A. C. Assessing the significance of focal activations using their spatial extent. Hum. Brain Mapp. 1, 210–220 (1994).

37. Braak, H., Alafuzoff, I., Arzberger, T., Kretzschmar, H. & Tredici, K. Staging of Alzheimer disease-associated neurofibrillary pathology using paraffin sections and immunocytochemistry. Acta Neuropathol. 112, 389–404 (2006).

38. Desgranges, B. et al. The neural substrates of memory systems impairment in Alzheimer’s disease. A PET study of resting brain glucose utilization. Brain 121 (Pt 4, 611–31 (1998).

39. Serrano-Pozo, A., Frosch, M. P., Masliah, E. & Hyman, B. T. Neuropathological Alterations in Alzheimer Disease. Cold Spring Harb. Perspect. Med. 1, a006189–a006189 (2011).

40. Kringelbach, M. L. & Rolls, E. T. The functional neuroanatomy of the human orbitofrontal cortex: evidence from neuroimaging and neuropsychology. Prog. Neurobiol. 72, 341–72 (2004).

41. Rolls, E. T. The brain and emotion. (Oxford University Press, 1999).

42. Grill-Spector, K., Kourtzi, Z. & Kanwisher, N. The lateral occipital complex and its role in object recognition. Vision Res. 41, 1409–1422 (2001).

43. Mamashli, F., Hämäläinen, M., Ahveninen, J., Kenet, T. & Khan, S. Permutation Statistics for Connectivity Analysis between Regions of Interest in EEG and MEG Data. Sci. Rep. 9, 7942 (2019).

44. Glasser, M. F. et al. A multi-modal parcellation of human cerebral cortex. Nature 536, 171–178 (2016).

45. Wang, H. et al. Functional Brain Connectivity Revealed by Sparse Coding of Large-Scale Local Field Potential Dynamics. Brain Topogr. 32, 255–270 (2019).

46. Nolte, G. et al. Identifying true brain interaction from EEG data using the imaginary part of coherency. Clin. Neurophysiol. 115, 2292–2307 (2004).

47. Zou, X., Coyle, D., Wong-Lin, K. F. & Maguire, L. Beta-amyloid induced changes in A-type K + current can alter hippocampo-septal network dynamics. J. Comput. Neurosci. 32, 465–477 (2012).

48. Styr, B. & Slutsky, I. Imbalance between firing homeostasis and synaptic plasticity drives early-phase Alzheimer’s disease. Nat. Neurosci. 21, 463–473 (2018).

49. Frere, S. & Slutsky, I. Alzheimer’s Disease: From Firing Instability to Homeostasis Network Collapse. Neuron 97, 32–58 (2018).

## References

50. Oldfield, R. C. The assessment and analysis of handedness: The Edinburgh inventory. Neuropsychologia 9, 97–113 (1971).

51. Rosen, W. G., Terry, R. D., Fuld, P. A., Katzman, R. & Peck, A. Pathological verification of ischemic score in differentiation of dementias. Ann. Neurol. 7, 486–488 (1980).

52. Reisberg, B., Ferris, S. H., de León, M. J. & Crook, T. The Global Deterioration Scale for assessment of primary degenerative dementia. Am. J. Psychiatry 139, 1136–1139 (1982).

53. Hollander, M., Wolfe, D. A. & Chicken, E. Nonparametric statistical methods (Vol. 751). (John Wiley & Sons, 2013).

