## Supplementary Material for "Cluster-permutation statistical analysis for high-dimensional brain-wide functional connectivity mapping"

Jose M. Sanchez-Bornot[1], Maria E. Lopez[2,3], Ricardo Bruña[2,3,4], Fernando Maestu[2,3,4], Vahab Youssofzadeh[5], Su Yang[1], Paula L. McLean[6], Girijesh Prasad[1] and KongFatt Wong-Lin[1]

[1] Intelligent Systems Research Centre, School of Computing, Engineering and Intelligent Systems, Ulster University, Magee campus, Derry~Londonderry, UK.

[2] Department of Experimental Psychology, Cognitive Processes and Speech Therapy Universidad Complutense de Madrid, Madrid, Spain.

[3] Networking Research Center on Bioengineering, Biomaterials and Nanomedicine, Madrid, Spain.

[4] Laboratory of Cognitive and Computational Neuroscience, Center for Biomedical Technology, Complutense University of Madrid and Technical University of Madrid, Madrid, Spain.

[5] Department of Neurology, Medical College of Wisconsin, Milwaukee, USA.

[6] Northern Ireland Centre for Stratified Medicine, Biomedical Sciences Research Institute, Ulster University, Northern Ireland, United Kingdom.

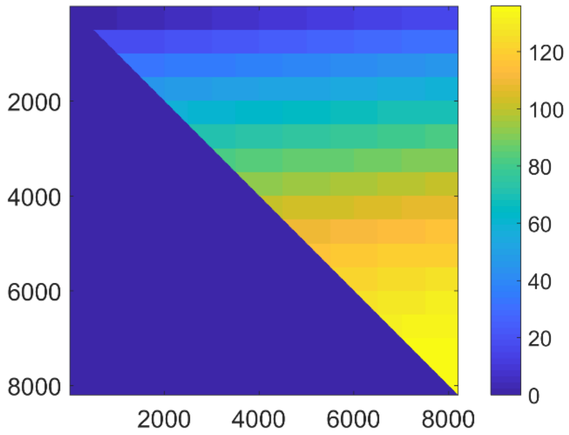


**Supplementary Figure 1: Block partition of 8196x8196 connectivity matrix into 16x16 sub-blocks.** The $C_2^{17} = 136$ blocks are arranged from left-to-right first then from-top-to-bottom in space. Only the block indices corresponding to the strictly upper triangular part are considered because of the symmetry of used FC measure.

**Supplementary Table 1: Participants' demographics**. The p-values were obtained by two-independent samples t-test ($*$) or chi-square test (+). HC = Healthy Control; MCI = Mild Cognitive Impairment; M = Male; F = Female; MMSE = Mini-Mental State Examination; IRM: Immediate Recall Memory; DRM: Delayed Recall Memory; RH_ICV and LH_ICV: right and left hippocampal volume normalized with intracranial volume, respectively. Highest education completed, using five levels: 1. Illiterate, 2. Primary studies, 3. Elemental studies, 4. High school studies, and 5. University studies.

| | **HC group (N=30)** | **MCI group (N=30)** | **p-values** |
|---|---|---|---|
| **Age (years)** | 72.1 ± 4.1 | 72.2 ± 4.0 | 0.874* |
| **Gender (M/F)** | 14/16 | 15/15 | $0.796^{+}$ |
| **Educational level** | 3.6 ± 1.1 | 3.4 ± 1.3 | 0.707* |
| **MMSE** | 29.2 ± 0.8 | 26.9 ± 1.9 | <1e-6* |
| **IRM** | 38.7 ± 8.0 | 18.9 ± 9.0 | <1e-8* |
| **DRM** | 24.6 ± 6.6 | 7.2 ± 8.0 | 0.089* |
| **RH_ICV** | 0.00250 ± 0.00030 | 0.00207 ± 0.00047 | 0.0002* |
| **LH_ICV** | 0.00251 ± 0.00035 | 0.00207 ± 0.00046 | 0.0002* |

**Supplementary Table 2: Algorithm for cluster parcellation of supra-threshold FC using breadth-first search.**

---

**Inputs:** $\{(X_{I_1(k)}, X_{I_2(k)}) : \{1, \dots, C_2^{8196}\} \vdash (\mathcal{R}^3, \mathcal{R}^3)\}$, which is an array of the supra-threshold FC links. By definition, $I_1(k) < I_2(k)$ as the FC matrix is symmetric and only the connection entries in the strictly upper triangular part are considered, where $I(k) = (I_1(k), I_2(k)) : \{1, \dots, C_2^{8196}\} \vdash \{1, \dots, 8195\} \times \{2, \dots, 8196\}$ is a functional mapping of the connection to its vertices index.

---

Insert the connection indices $(I_1(k), I_2(k))$, $k \in \{1, \dots, C_2^{8196}\}$, in a doubly-linked list $DLink$.<br>
While $DLink$ is not empty<br>
    Set an empty singly-linked list $SLink$ and an empty $Cluster$;<br>
    Extract a connection from the top of $DLink$.<br>
    Insert it in $SLink$ and assign it to the current $Cluster$;<br>
    While $SLink$ is not empty<br>
        Extract the connection $(I_1(k), I_2(k))$ from the top of $SLink$;<br>
        For each connection $(I_1(l), I_2(l))$ in $DLink$<br>
            Check whether $I_m(k)$ is equal to $I_n(l)$ for some $m, n \in \{1,2\}$;<br>
            If true<br>
                set $J(k) = I_{3-m}(k)$ and $J(l) = I_{3-n}(l)$;<br>
            End If<br>
            Check whether vertices $X_{J(k)}$ and $X_{J(l)}$ are neighbours in the cortical surface;<br>
            If true<br>
                Insert the connection $(I_1(l), I_2(l))$ at the bottom of $SLink$;<br>
                Insert it into the current $Cluster$;<br>
                Remove it from $DLink$;<br>
            End If<br>
        End For<br>
    End While<br>
End While

---

**Supplementary Table 3: MATLAB custom code for an efficient rank computation within the permutation approach.** See the implementation of "mysortedtiedrank" function in Supplementary Table 4:

```
[MA,N] = size(XA); % XA: MCI data. Size: MAxN

MB = size(XB,1); % XB: HC data. Size: MBxN

M = MA + MB;

X = [XA; XB];

N = 1000; % Monte-Carlo randomizations

RS = zeros(1,N); % rank-sum stat

rank = zeros(1,M);

[Xsort,indsort] = sort(X); % sorting per column

for it = 1:N

    indperm = randperm(M);

    rsort = mysortedtiedrank(Xsort(:,it));

    rank(indsort(:,it)) = rsort(:,it);

    for k = 1:MA

        RS(it) = RS(it) + rank(indperm(k));

    end

end
```

**Supplementary Table 4: MATLAB custom code of "mysortedtiedrank" function.**

```
function rs = mysortedtiedrank(xs)

M = length(xs);

up = zeros(1,M); % to keep position for unique elements and its number of repetitions

uc = zeros(1,M); % to keep number of repetitions for unique elements

ui = zeros(1,M); % to keep initial position for unique elements

up(1) = 1;

ui(1) = 1;

indcurr = 1;
```

```
for it = 2:M

    if (xs(it) == xs(it-1))

        up(1,it) = indcurr;

        uc(indcurr) = uc(indcurr) + 1;

        ui(it) = ui(it-1);

    else

        indcurr = indcurr + 1;

        up(1,it) = indcurr;

        ui(it) = it;

    end

end

tmp = 0.5*uc(1:indcurr);

rs = tmp(up) + ui;
```

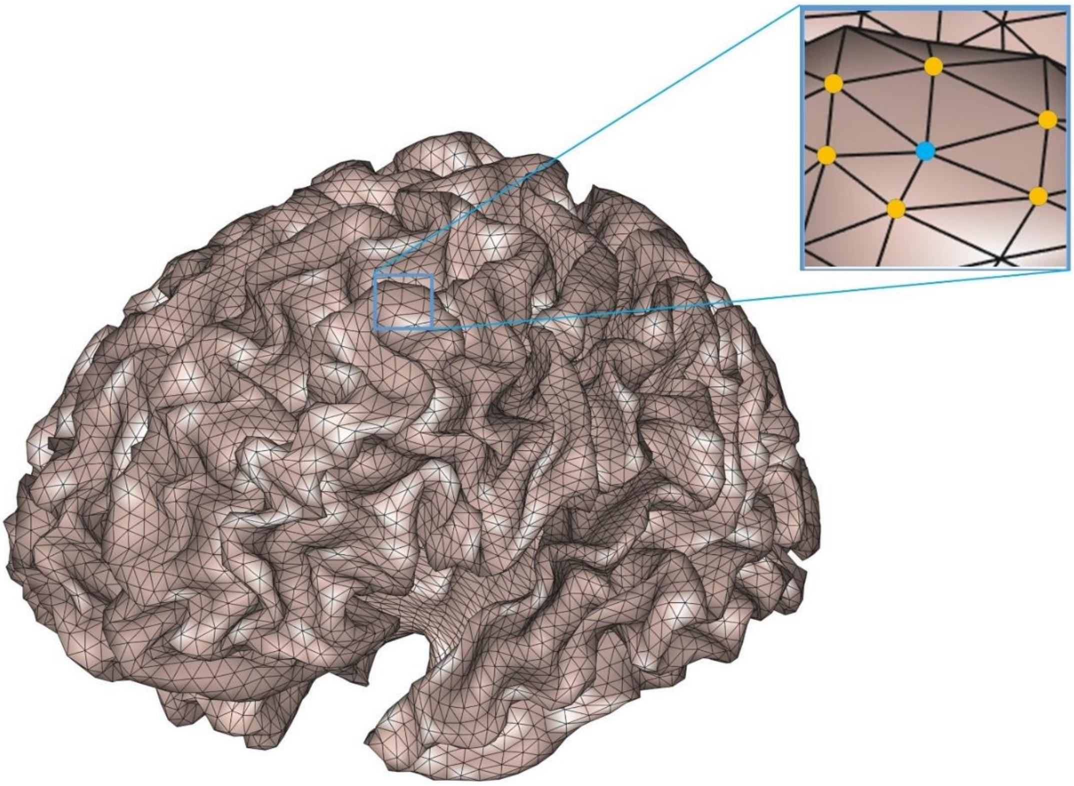

**Supplementary Figure 2: Mesh of the cortical surface for the SPM12 medium template of 8196 vertices.** The inset shows the adopted "neighbour" relationship between any vertex (highlighted in blue colour) with its surrounding vertices (orange colour). In general, two vertices are considered neighbours in the cortical surface if they share an edge of the mesh.

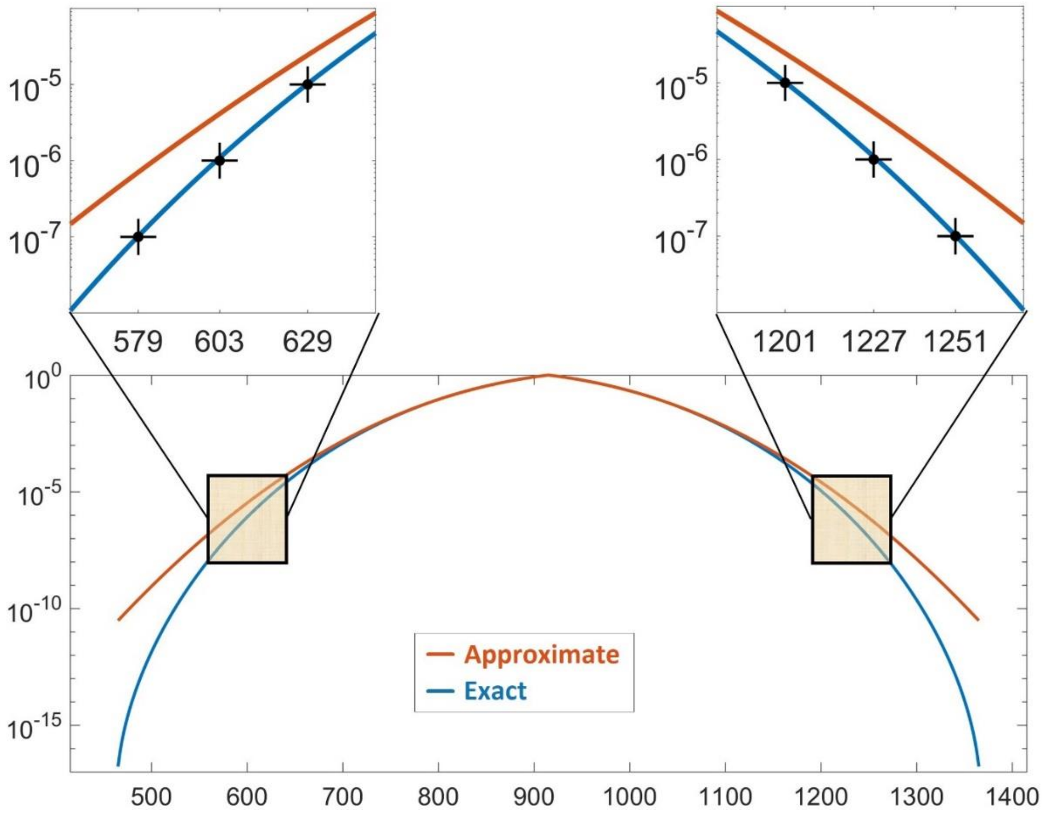


**Supplementary Figure 3: Plot of the approximate (orange colour) and exact statistic (blue colour) for the Wilcoxon rank-sum analysis between measurements of 30 MCI and 30 HC participants.** The insets show the *a priori* selected supra-threshold values with its corresponding statistic critical values for the lower and upper tails, respectively.

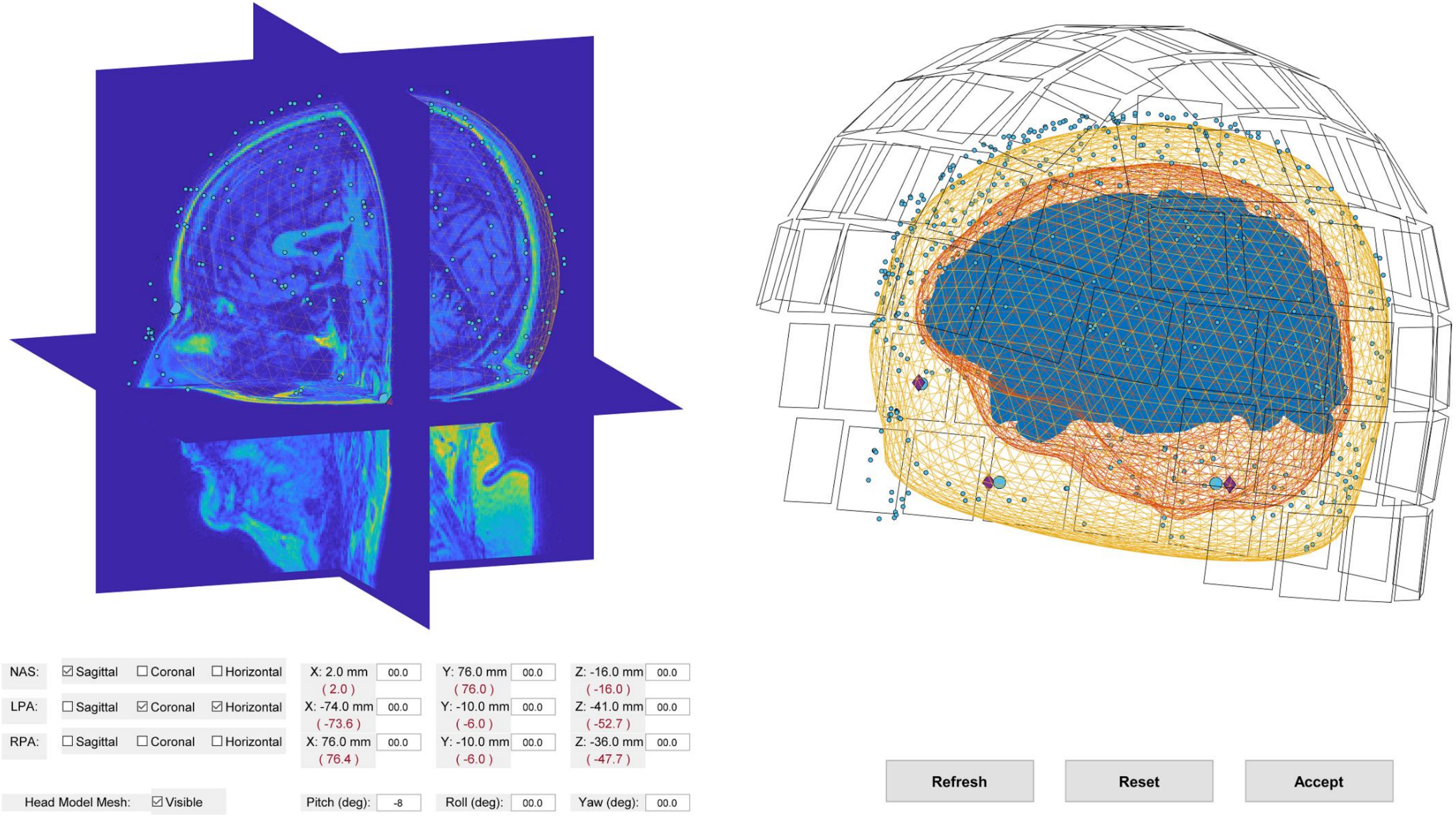


**Supplementary Figure 4: Graphical user interface of custom co-registration routine based on SPM12 toolbox.**
